# HDAC11 is a fatty-acid deacylase

**DOI:** 10.1101/212043

**Authors:** Zsofia Kutil, Zora Novakova, Marat Meleshin, Jana Mikesova, Mike Schutkowski, Cyril Barinka

## Abstract

Histone deacetylase 11 (HDAC11) is a sole member of the class IV HDAC subfamily with negligible intrinsic deacetylation activity. Here we report *in vitro* profiling of HDAC11 deacylase activities, and our data unequivocally show that the enzyme efficiently removes acyl moieties spanning 8–18 carbons from the side chain nitrogen of the lysine residue of a peptidic substrate. Additionally, N-linked lipoic acid and biotin are removed by the enzyme, although with lower efficacy. Catalytic efficiencies toward dodecanoylated and myristoylated peptides exceed 70,000 M^−1^s^−1^ making HDAC11 the most proficient fatty acid deacylase of the HDAC family. Interestingly, HDAC11 is strongly inhibited by free myristic, palmitic and stearic acids with inhibition constants of 6.5 µM, 0.9 µM, and 1.6 µM, respectively. At the same time, its deacylase activity is stimulated more than 2.5-fold by both palmitoyl-coenzyme A and myristoyl-coenzyme A, pointing toward metabolic control of the enzymatic activity by fatty acid metabolites. Our data reveal novel enzymatic activity of HDAC11 that can, in turn, facilitate the uncovering of additional biological functions of the enzyme as well as the design of isoform-specific HDAC inhibitors.

---

Dynamic and reversible acetylation at the ε-NH_2_ group of lysine has emerged as a critical regulatory mechanism in living organisms. It is implicated in signaling cascades, protein-protein interactions, localization and degradation, chromatin assembly, DNA repair and recombination, and metabolic stress response. Moreover, owing to the widespread use of high-resolution mass spectrometry, other types of acyl modifications, including formylation, propionylation, butyrylation, crotonylation, malonylation, succinylation, glutarylation, lipoylation, and myristoylation, have been recently identified expanding thus the portfolio of post-translation modifications (PTMs) that control a number of cellular processes.^1–3^

Acylation status of proteins is defined by counter-balancing activities of histone acetyltransferases (HATs) and histone deacetylases (HDACs).^4,5^ Additionally, non-enzymatic acylations by metabolic intermediates/end products (e.g. acyl-coenzyme A; acyl-CoA) were also described, linking thus lysine acylation directly to the metabolic state of a given cell/organelle.^6–9^ On the contrary, the reverse reaction, i.e. the removal of acyl groups from the lysine side chain, is more tightly regulated by substrate specificities of individual HDACs and their spatiotemporal distribution within the cell.

HDACs are evolutionarily conserved among organisms and can be divided into 4 classes based on sequence homology and enzymatic mechanism. Members of class I (HDAC 1, 2, 3, and 8), II (HDAC 4 – 7, 9 and 10), and IV (HDAC11) are zinc dependent hydrolases, while class III proteins (called sirtuins; SIRT 1 – 7) represent a distinct group encompassing NAD^+^ dependent enzymes.^5^ HDAC11 is the most recently identified sole member of the class IV HDAC subfamily, and its functional role in mammalian (patho)physiology remains poorly understood.^10^ At the protein level, the sequence analysis of the human enzyme revealed high conservation of critical residues in the catalytic core regions shared by both class I and II mammalian HDAC pointing toward catalytic proficiency of HDAC11. At the same time, however, very low deacetylation activity has been ascribed to the protein, and there is no direct evidence linking HDAC11 deacetylase activity to observed physiology effects, such as its involvement the immune system regulation or modulation of cancer growth.^11–14^

## Results and Discussion

To facilitate understanding of the biological role(s) of HDAC11, we set out to map the substrate specificity of HDAC11 *in vitro*. To this end, we first cloned HDAC11 from *H. sapiens* (hsHDAC11, NP_079103.2, amino acids 1 – 347) and *D. rerio* (drHDAC11, NP_001002171.1, amino acids 1 – 366) and expressed both proteins using the HEK293/T17-based heterologous system. Additionally, putative catalytically active histidine residues were mutated to alanines (H142A, H143A; hsHDAC11 numbering) to prepare enzymatically impaired mutants for negative control experiments. All recombinant proteins were purified to near homogeneity by Streptactin-affinity chromatography and were used for the ensuing biochemical studies (Fig. 1).

**Figure 1:**
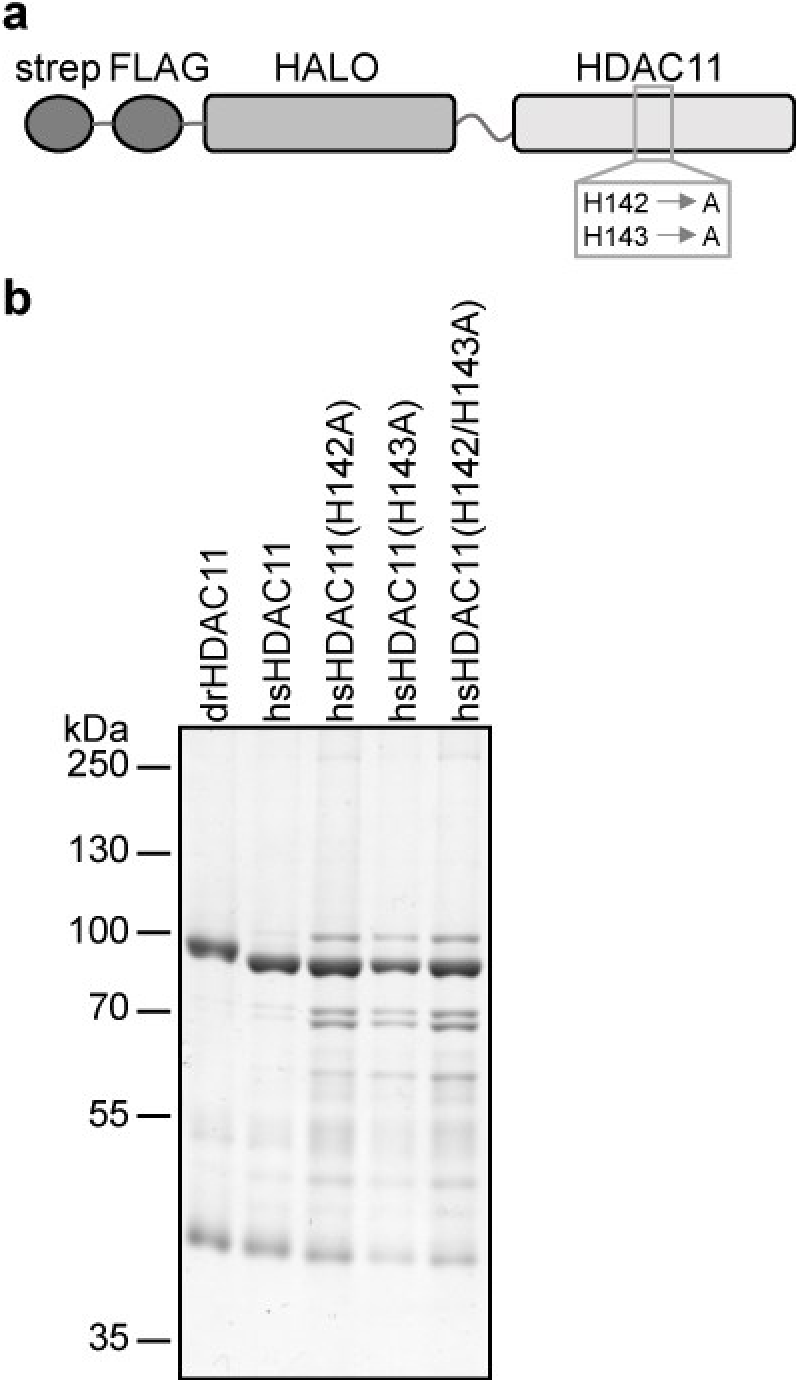
Expression and purification of HDAC11 variants. drHDAC11 and hsHDAC11, along with catalytically impaired H142A/H143A mutants, were cloned as fusions with the N-terminal Strep-FLAG-HALO tag. Fusion proteins were expressed in HEK293/T17 cells and purified using Streptactin affinity chromatography. **A**: Schematic representation of constructs used in this study. **B**: Coomassie-stained reducing SDS-PAGE analysis of purified HDAC11 constructs. The major contaminating species include α, β-tubulin, heat-shock proteins, and the cleaved Strep-FLAG-HALO fusion partner.

Both hsHDAC11 and drHDAC11 were profiled against a panel of acyl derivatives of Abz-Ser-Arg-Gly-Gly-[Lys-Acyl]-Phe-Phe-Arg-Arg-NHM_2_, a synthetic nonapeptide (Fig. 2A). The peptide sequence was designed based on peptide microarray screening experiments and is efficiently recognized by both class I and class II HDACs (data not shown). As no physiologically relevant HDAC11 substrates have been identified thus far, the use of such an “artificial” peptide sequence is fully warranted for *in vitro* profiling and comparable to, for example, the histone-derived H3K9 sequence, provided it is recognized by tested enzymes.

**Figure 2:**
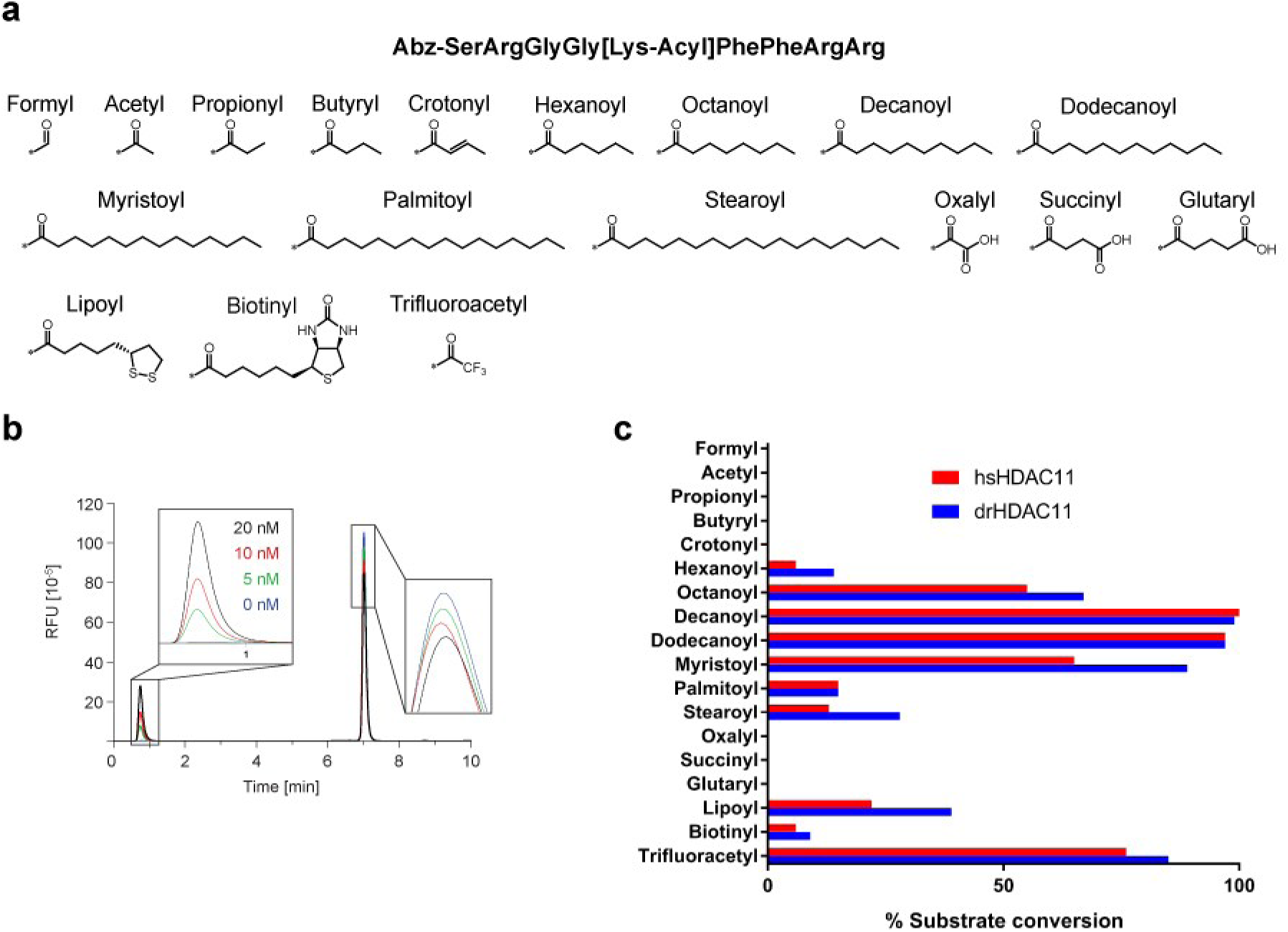
Deacylation profiling of HDAC11 substrates. **A:** Sequence of the synthetic fluorescent peptide substrate together with the scheme of acyl derivatives used in this study. Abz – fluorescent aminobenzoyl group used for substrate/product quantification. **B:** Example of an HPLC profile used to quantify deacylation reactions based on Abz fluorescence. Varying concentrations of HDAC11 (0, 5, 10, and 20 nM) were incubated with 100 µM Abz-SRGG[K-Myr]FFRR-NH_2_ in the HEPES buffer and deacylation reactions carried out for 30 mins. at 37°C. **C**: Percentage of an acylated substrate converted to the deacylated product in 30 mins at 37°C using 100 nM HDAC11 and 10 µM substrate.

In total, 18 peptide-derivatives with different acyl modifications were synthesized, and the purity/identity of the resulting side chain-acylated peptides were confirmed using ultra-high performance liquid chromatography coupled with single quadrupole mass spectrometry (Figs. 2A and S1-S17). For the substrate profiling, 100 nM HDAC11 was mixed with 10 µM substrate in an HEPES buffer. Deacylation reactions were carried out for 30 mins at 37°C, and the extent of deacylation was quantified using HPLC (Fig. 2B). HDAC11 was inactive against formylated, acetylated, propionylated, butyrylated, and crotonylated peptides as well as peptides carrying negatively charged acyl residues (oxalyl, succinyl, glutaryl). However, surprisingly, both hsHDAC11 and drHDAC11 efficiently removed acyl groups spanning 6-18 carbons from the peptide sequence, revealing that HDAC11 can act as a proficient fatty-acid deacylase. Under assay conditions, 3% - 100% of medium-to-long fatty-acylated substrates were converted to the corresponding product, with decanoyl and dodecanoyl being the most efficient substrates with conversions close to 100%. Furthermore, HDAC11 also removes two additional physiologic modifications – lipoyl and biotinyl – with conversions of 22% and 6%, respectively. The non-biologically relevant trifluoroacetyl (TFA) group, which is used as a super-substrate for the class IIa HDACs, was removed by HDAC11 with 71% conversion. It is interesting to note that acyl preferences of both human and zebra fish proteins are virtually identical, indicating functional phylogenetic conservation of HDAC11 acyl specificities.

In order to gain more quantitative insight into hsHDAC11 acyl preferences, we carried out steady state kinetic analyses of selected acyl-peptides, and the results are summarized in Table 1 and the Supplementary Figure S18. The decanoyl, dodecanoyl and myristoyl groups were the most efficiently removed acyl residues with overall catalytic efficiencies of (k_cat_/K_M_) 77,722 s^−1^M^−1^, 149,000 s^−1^M^−1^, and 72,143 s^−1^M^−1^, respectively. For the longer acyl chains (16 – 18 carbons), the catalytic efficiency decreased dramatically primarily due to K_M_ values that were more than 100 times higher (> 500 µM). Clearly, within the context of a peptidic substrate, both palmitoyl/stearoyl moieties are too long to be efficiently accommodated by the enzyme. For the hexanoylated peptide, the shortest acyl group removed efficiently by HDAC11 under our experimental conditions, there is approximately a 10-fold increase in the K_M_ value (68 µM) with the concomitant, approximately 20-fold decrease in the corresponding reaction rate.

**Table 1:** Steady-state kinetics of deacylation of selected substrates by hsHDAC11. Substrates were used in concentrations of 0.01 – 200 µM. Michaelis-Menten constants (K_M_ and *k*_*cat*_) for individual peptides along with the catalytic efficiency were calculated from the non-linear regression fit using the GraphPad program. Data represent mean values ± s.d. (n = 3).

| Acyl group | $K_M$ [$\mu\text{M}$] | $k_{cat}$ [$\text{s}^{-1}$] | $k_{cat}/K_M[\text{s}^{-1}\text{M}^{-1}]$ |
| --- | --- | --- | --- |
| Hexanoyl | $68.0 \pm 10.6$ | $0.019 \pm 0.001$ | 279 |
| Octanoyl | $13.3 \pm 2.1$ | $0.047 \pm 0.002$ | 3,534 |
| Decanoyl | $7.9 \pm 3.1$ | $0.610 \pm 0.060$ | 77,722 |
| Dodecanoyl | $2.0 \pm 0.5$ | $0.290 \pm 0.020$ | 149,000 |
| Myristoyl | $1.4 \pm 0.3$ | $0.101 \pm 0.005$ | 72,143 |
| Palmitoyl | >500 | >0.060 | ND |
| Stearoyl | >500 | >0.080 | ND |
| Biotin | >200 | >0.030 | ND |
| Lipoyl | $10.6 \pm 2.1$ | $0.025 \pm 0.001$ | 2,358 |
| Trifluoroacetyl | >200 | >0.700 | ND |

Interestingly, there is a marked difference between the kinetics of delipoylation and debiotinylation reactions. While the lipoyl moiety is recognized and removed by HDAC11 quite efficiently with respective kinetic constants K_M_ = 10.6 µM and k_cat_ = 0.025s^−1^, we could not accurately determine kinetic constants for the biotinylated peptide as only the linear part of the Michaelis-Menten plot was sampled up to 200 µM substrate concentration (Fig. S18). Compared with kinetic data on fatty acylated substrates, the lipoyl moiety is removed with efficacy similar to that of the octanoyl group, its biological precursor. In the case of the bulkier and more polar biotinyl group, the slower substrate turnover is linked to both a decreased reaction rate and increased K_M_ values.

SIRTs 1-3 and SIRT6 displayed deacylase activities against H3K9Myr and H3K9Dodeca peptides with reaction rates of 0.01 – 0.1 s^−1^.^15^ Similarly, HDAC8 is able to deacylate H3K9 FA-modified peptide at the rate of 0.0005 s^−1^ (depalmitoylation) to 0.02 s^−1^ (deoctanoylation).^16^ Compared with these data, HDAC11 removes acyl groups more robustly, with deacylation rates up to 0.60 s^−1^ and catalytic efficiencies >150,000 M^−1^s^−1^. Nevertheless, such comparisons shall be interpreted with caution. First, neither H3K9 nor our synthetic peptide represents true physiological substrates of the enzymes, and different peptidic sequences could be deacylated at markedly different rates. Secondly, differences in enzymatic preparations (*E.coli* vs baculovirus vs HEK293T heterologous expressions) and/or the presence/absence of posttranslational modifications shall be taken into account. Finally, peptides are only surrogates for physiological substrates, and a deacylation rate of natively folded proteins can be orders of magnitude higher compared with corresponding peptide sequences.^17^ Nevertheless, given the superior catalytic efficiency of hsHDAC11 (as compared with other HDACs/SIRTs), it is clear that HDAC11 is by far the most proficient fatty acid (FA) deacylase reported to date underscoring thus the physiological relevance of our findings.

To ascertain that observed deacylase activity does not result from possible contamination by other HDACs/SIRTs^15, 16^ ^18–22^, we first analyzed our preparations using mass spectrometry. The high-resolution LC-MS/MS data revealed that the major contaminating species in our preparations include “common contaminants,” such as αβ-tubulin and heat shock proteins with virtually absent HDACs/SIRTs (<0.001%; Supplementary table S2). Moreover, we prepared three HDAC11 mutants with impaired hydrolytic activity, namely hsHDAC11(H142A), hsHDAC11(H143A), and hsHDAC11(H142A/143A). The mutations were introduced based on the amino acid sequence alignment of Zn^2+^-dependent HDACs, where one (or both) of the histidine residues are required for their hydrolytic activity. Mutated proteins were assayed using protocols identical to wild-type enzyme preparation with myristoylated peptide as a substrate. Interestingly, while the hsHDAC11(H142A) and hsHDAC11(H143A) retained approximately 7% and 0.3% activity of the wild type hsHDAC11, respectively, the hsHDAC11(H142A/143A) construct was fully inactive (Fig. S19). Our mutagenesis data thus confirm that observed fatty-acid deacylase activity is indeed the intrinsic catalytic property of HDAC11. Additionally, both histidine residues seem to be critical for the HDAC11 activity, with H143 likely acting as the general base, while H142 served as a general electrostatic catalyst similar to corresponding residues inHDAC8.^23^

Deacetylase activity of SIRT6 was reported to be stimulated by free long chain FAs.^15^ To analyze the effect of free FAs on deacetylase activity of HDAC11, the enzyme was preincubated with 100 µM FAs for 15 mins at 37°C and its activity determined using the trifluoroacetylated peptide substrate. Contrary to the reported FA-dependent stimulation of SIRT6 deacetylase activity, we observed strong (more than 50%) inhibition of hsHDAC11 by free decanoic through stearic acid (Fig. 3A). Inhibition curves were generated for FAs positive in the screening, and the results are shown in Fig. 3B. Palmitic acid inhibits hsHDAC11 the most potently with an IC_50_ value of 900 nM, followed by stearic acid (IC_50_ = 1.2 µM) and myristic acid (IC_50_ = 6.9 µM). On one hand, HDAC11 inhibition by free FAs corroborates our findings that the enzyme preferentially recognizes longer alkyl chains and affinity decreases with decreasing length of the acyl group. Hexanoylated peptide and free hexanoic acid represent a very poor substrate and inhibitor, respectively, and shorter fatty acids are neither inhibitory nor their peptide forms can serve as HDAC11 substrates. At the same time, however, there is a clear difference in interactions between HDAC11 and free FAs on one side or acyl groups in the context of peptidic substrates on the other side. In the case of free FAs, palmitic and stearic acids interact with the enzyme most avidly, while corresponding acyl groups in the context of peptidic substrates are less preferred as compared with myristoyl/dodecanoyl groups.

**Figure 3:**
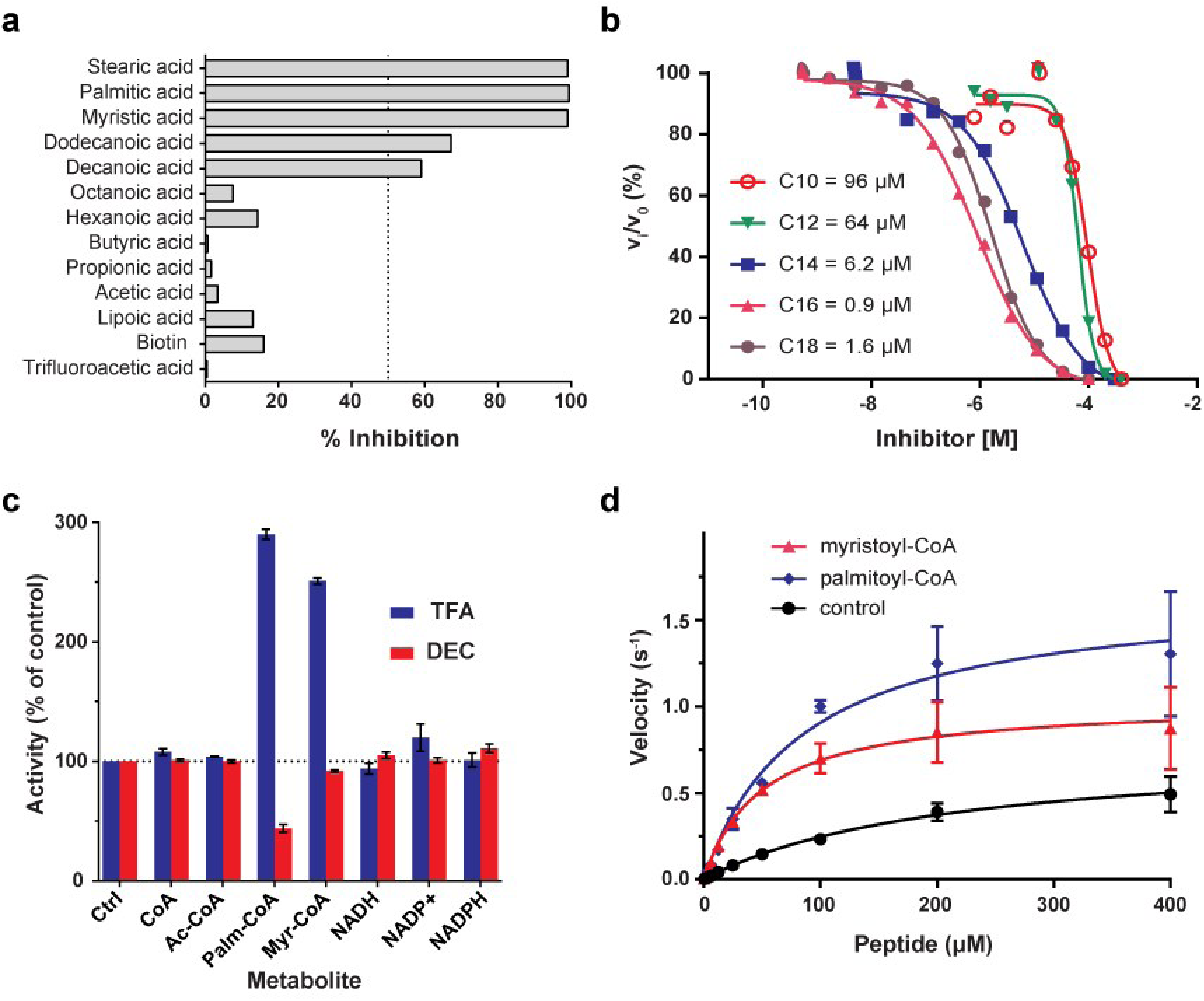
Modulation of HDAC11 activity by free acids and intermediate metabolites. **A:** Inhibitory activity of free acyl groups at 100 µM concentration. 10 nM HDAC11 was preincubated with 100 µM concentration of a given compound for 10 mins at 37°C. The deacylation reaction was started by the addition of 10 µM Abz-SRGG[K-TFA]FFRR-NH_2_, terminated after 30 mins and analyzed by HPLC. Data are shown as the fraction of inhibited vs control (without any compound) reactions. **B:** Full-dose response inhibition curves for compounds with higher than 50% inhibitory activity under screening conditions. Corresponding IC_50_ values were calculated from the inhibition curves using the GraphPad Prism software (GraphPad Software, San Diego, CA, USA). Data are plotted as mean values ± s.d. from three independent experiments (n = 3). **C:** 10 nM HDAC11 was preincubated with 100 µM concentration of a given metabolic intermediate and its activity against 10 µM trifluoroacetylated (TFA; blue) or decanoylated (DEC, red) substrates was analyzed as in panel A. While there is a limited influence on HDAC11 activity of majority of intermediated, palmitoyl-CoA/myristoyl-CoA activate/inhibit HDAC11 in a substrate-dependent manner. **D:** Steady-state kinetics of HDAC11 toward Abz-SRGG[K-TFA]FFRR-NH_2_ in the presence of 100 µM palmitoyl-CoA (blue diamonds) and myristoyl-CoA (red triangles). An increase in the HDAC11 catalytic efficiency over the control reaction (black squares) results from approximately a 10-fold decrease in the K_M_ value.

Upon entry into the cell, free fatty acids of >12 carbon chains are rapidly conjugated to coenzyme A (CoA) forming acyl-CoA adducts in the cytoplasm.^24^ We thus tested the effect of palmitoyl-CoA and myristoyl-CoA on HDAC11 activity. Contrary to free FAs, palmitoyl-CoA and myristoyl-CoA (100 µM) stimulate HDAC11 deacetylation activity by 2.9-fold and 2.5-fold, respectively, using the trifluoroacetylated peptide substrate. Concurrently, however, palmitoyl-CoA inhibits the removal of decanoyl moiety from the substrate peptide by more than 70% (Fig. 3C). As observed for other family members, HDAC allosteric modulation by metabolic intermediates is clearly substrate dependent, and our additional analysis revealed that the presence of acyl-CoAs increased HDAC11 activity by decreasing its K_M_ value for the trifluoroacetic substrate (Fig. 3D).^25, 26^ Additionally, we tested the direct regulation of HDAC11 activity by other metabolic intermediates, including free CoA, acetyl-CoA, NAD^+^, NADPH, and NADP^+^. HDAC11 was incubated with decanoylated or trifluoroacetylated substrates in the presence of the metabolite (100µM) and HDAC11 deacylase activity analyzed using an HPLC-based assay. No marked influence on HDAC11 enzymatic activity was detectable, revealing that the FA moiety is essential for effective HDAC11 interactions.

Modulation of HDAC11 activity by physiological concentrations of free FAs and their metabolic intermediates suggests that the enzyme can serve as a sensor for fatty acid metabolism in the cell. HDAC11 could arguably deacylate proteins that would be modified non-enzymatically by reactive FA-CoAs that are formed as intermediates of fatty acid synthesis pathway.^8^ HDAC11 could also function as a binding/transport protein for free FAs/acyl-CoAs. Furthermore, it would be interesting to see whether HDAC11 could play other roles in fatty acid metabolism, for example by hydrolyzing small acylated molecules (metabolites, metabolic intermediates) similarly to preferential deacetylation of small acetylated polyamines by HDAC10.^27^ Finally, we believe that the inhibition of HDAC11 by FAs can serve as a starting point for the design of HDAC11-specific inhibitors used as tools for biological studies.

At present, a limited number of proteins fatty acylated at the N^ε^ lysine group has been identified, with the myristoylated TNFα being the most prominent example.^20^ We thus used the myristoylated TNFα-derived peptide to assess demyristoylase activity of HDAC11 in “a more physiologically relevant model.” Our data show that the peptide is efficiently demyristoylated by HDAC11 with K_M_ and k_cat_ values of 0.73 µM and 0.027 s^−1^, respectively (Fig. S18). These findings thus corroborate and extend our *in vitro* screening results. Furthermore, similar K_M_ values (as approximations of the substrate affinity), for both the library-derived and TNFα-derived peptides, suggest that the acyl chain rather than the peptide sequence *per se* can be the driving force of the high affinity of myristoylated substrates.

The three-dimensional structure of HDAC11 has not been reported to date. To elucidate the putative binding mode of a decanoylated tripeptide in the substrate-binding cavity of HDAC11, we used the Modeller 9.14 software to construct an hsHDAC11 homology model based on the HDAC8 X-ray structure (3MZ4) as a template. Next, we docked a Gly-[Lys-decanoyl]-Phe tripeptide (derived from the tested substrate) into the hsHDAC11 model using the AutoDock Vina 1.1.2. (Fig. 4). A narrow pocket, which accommodates the lysine side chain of the substrate, leads from the protein exterior to the Zn^2+^-dependent catalytic site, where it splits into two internal tunnels, lateral and vertical, which can in principle accommodate long aliphatic fatty acid chains of substrates. Our docking experiments suggest that the lateral tunnel, the entrance of which is delimited by the Lys33 and Phe141, is more likely to be occupied by acyl moieties of substrates (Fig. 4B).This tunnel, also known as the “foot pocket” in HDAC8, is believed to be an exit route for the free acetate in class I HDACs.^28, 29^ At the same time, it is missing in the class II HDACs apparently due to the presence of a salt bridge between Glu502 and Arg606 (human HDAC6 numbering) conjoining loops 1 and 3 of the class II HDACs.^30^ In this region, HDAC11 is highly similar to HDAC8 (and class I HDACs as an extension) because the corresponding glutamate and arginine residues are missing. This fact enables the formation of a deep lateral pocket in the HDAC11 structure. Consequently, we hypothesize that HDAC11 utilizes a catalytic mechanism similar to that of HDAC8, the only other Zn^2+^-dependent HDAC reported to remove longer (> 8 carbon atoms) acyl groups.^16^ The comparison of internal pockets of class I and II HDACs is clearly documented in Fig. 4, where the Gly-[Lys-decanoyl]-Phe tripeptide from our docking experiment is inserted into the superimposed models of HDACs 6, 8, and 11. While both HDAC8 and 11 binding pockets are spacious enough to accommodate the decanoyl moiety of the substrate, the internal cavity in HDAC6 is too small to harbor bulkier acyl groups.

**Figure 4:**
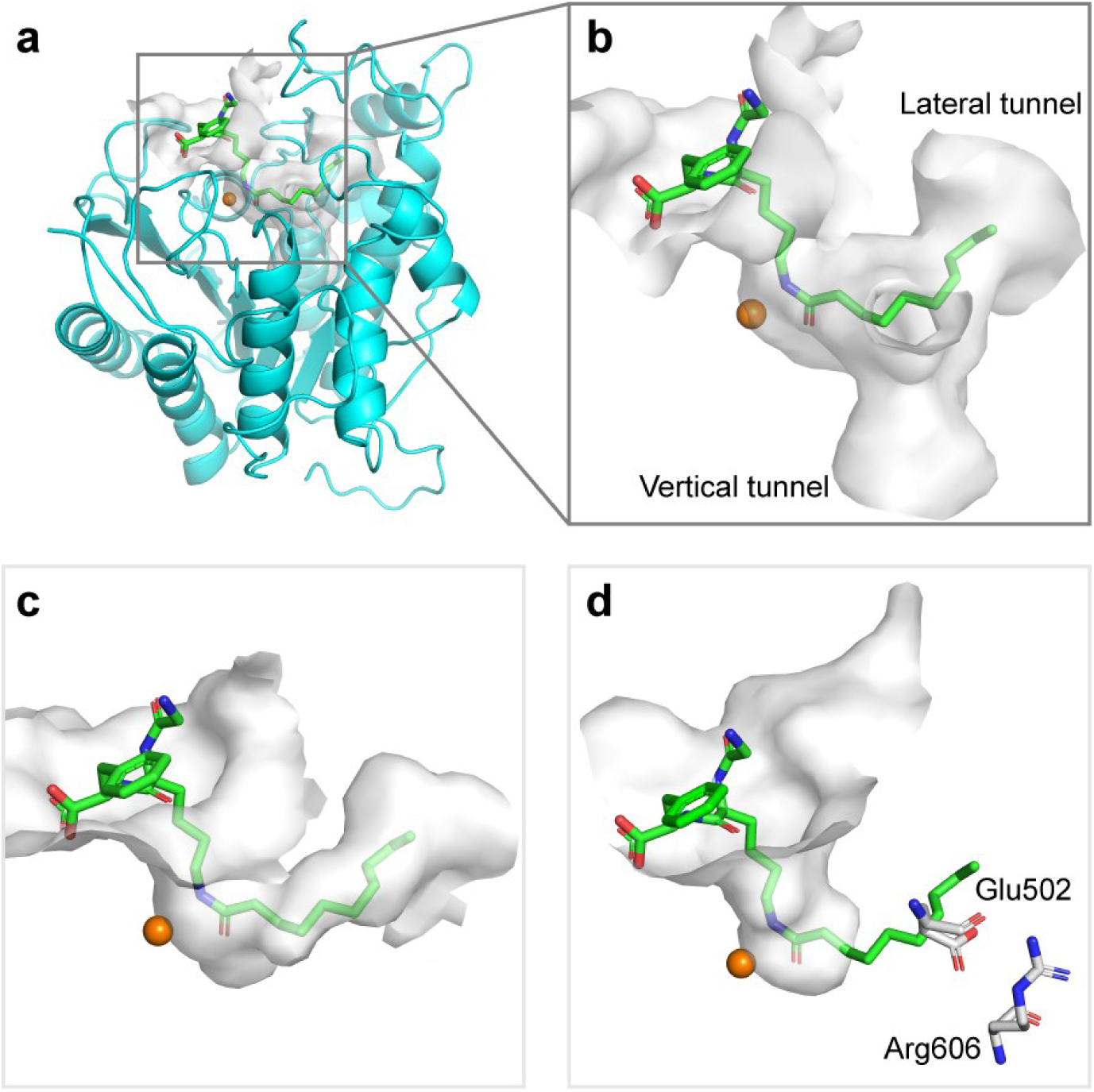
Homology modeling and docking. **A:** Gly-[K-Dec]-Phe tripeptide (green; stick representation) docked into the homology model of hsHDAC11 (cyan; cartoon representation). The 3D homology model of hsHDAC11 was constructed with the HDAC8 crystal structure (pdb code: 3MZ4) as a template using the Modeller 9.14. The docking experiment was carried out using AutoDock Vina 1.1.2. **B:** The detail of the hsHDAC11 internal pocket (surface shown as gray solid) with the Gly-[K-Dec]-Phe tripeptide docked. The active-site zinc ion is shown as an orange sphere. **C, D:** The docking pose captured by the tripeptide in the hsHDAC11 homology model was superimposed onto the crystal structure of HDAC8 (C; pdb code: 2V5W) and HDAC6 (D; pdb code: 5G0H). The comparison of HDAC11, 8, and 6 internal pockets reveals that while both HDAC8 and 11 internal pockets are sufficiently large to accommodate the decanoyl moiety of the substrate, the internal cavity of HDAC6 is too small to harbor longer alkyl groups. Glu502 and Arg606, which are responsible for the absence of the lateral tunnel in HDAC6, are shown in stick representation with carbons colored gray.

In conclusion, our data reveal for the first time that HDAC11 is the most proficient de-fatty acylase with preferences for longer alkyl chains (decanoyl/dodecanoyl) with catalytic efficiencies exceeding 150,000 M^−1^s^−1^. We also demonstrate that the HDAC11 enzymatic activity is modulated by free fatty acids and their metabolites. Beyond their potential biological ramifications, our results can be exploited as a starting point for the design of HDAC11-specific inhibitors for applications evaluating HDAC11 role(s) in cellular processes.

## METHODS

### Chemicals and peptides

If not stated otherwise, all chemicals were purchased from Sigma-Aldrich (St. Louis, MO, USA).

### HDAC11 cloning and mutagenesis

The codon-optimized DNA template was used for cloning of zebra fish HDAC11 (drHDAC11; NP_001002171.1). HDAC11 were custom-made by the Thermo Fisher gene-string synthesis protocol. The human HDAC11 (hsHDAC11; NP_079103.2) coding sequence was kindly provided by Prof. E. Seto (The George Washington University, D.C., USA). Coding sequences were PCR amplified with corresponding sets of gene-specific primers (Supplementary Table S1) and recombined using Gateway cloning protocols (Invitrogen, Carlsbad, CA, USA) to construct desired expression vectors as described previously.^17^ The expression clone for hsHDAC11 was further used as a template to generate catalytically impaired H142A/H143A mutants using the Quick-change site-directed mutagenesis approach. To this end, histidines 142 and 143 were mutated to alanines via PCR with mutagenic primers (Supplementary Table S1) followed by elimination of template by the DpnI treatment. The identity of all HDAC11 expression clones was verified using Sanger sequencing.

### HDAC11 expression and purification

All HDAC11 variants were expressed using HEK-293/T17 cells following transient transfection mediated by linear polyethylene imine (PEI; Polysciences Inc., Warrington, PA, USA) as described previously.^17^ Three days after transfection cells were harvested by centrifugation at 500xg for 10 mins and suspended in a lysis buffer (50 mM Tris, 150 mM NaCl, 10 mM KCl, 2 mM MgCl_2_, 10% glycerol, pH 8) supplemented with benzonase (2 U/ml; Merck, Darmstadt, Germany) and a cocktail of protease inhibitors (Roche, Basel, Switzerland). Cell lysis was enhanced by the addition of Igepal-630 (final concentration 0.2%) followed by incubation for 30 mins at 4°C. Cell lysate was cleared by centrifugation at 40,000xg for 30 mins at 4°C, and the supernatant was loaded on a Strep-Tactin column (IBA, Gottingen, Germany) previously equilibrated in the lysis buffer. The column was first washed with the lysis buffer supplemented with 2 mM ATP and 10 mM MgSO_4_ followed by the second wash with the elution buffer (50 mM HEPES, 100 mM NaCl, 50 mM KCl, 10% glycerol, pH 7.5). Fusion proteins were eluted with the elution buffer supplemented with 3 mM desthiobiotin. Eluted proteins were concentrated to 2 mg/ml and flash frozen in liquid nitrogen.

### Peptide synthesis and characterization

Details of the peptide synthesis and characterization are described in the accompanying Supplementary Material (Supplementary methods + Supplementary figures S1 – S17). Briefly, the peptides were synthesized using Fmoc-based solid-phase peptide synthesis (SPPS) with automated microwave peptide synthesizer Liberty Blue^TM^ (CEM Corporation, Matthews, NC, USA). The coupling of amino acids was performed with DIC/OxymaPure at 90 °C for 2 mins. Fmoc deprotection was accomplished with 20% piperidine solution in DMF at 90 °C for 1 min.

The lysine acylation was done either on the resin (formylation, oxalylation, succinylation, glutarylation, propionylation, butyrylation, crotonylation, trifluoroacetylation, and myristoylation) or in solution (hexanoylation, octanoylation, decanoylation, dodecanoylation, palmitoylation, stearoylation, lipoylation, and biotinylation). For the on-resin modifications, lysine was used in SPPS as a nosyl-protected derivative (Fmoc-Lys(Ns)-OH)^31^ and after peptide synthesis, the nosyl protecting group was selectively removed, and the respective free lysine side chain was modified according to the detailed procedure described in the Supplementary material. The peptide was then cleaved from the resin with TFA/H_2_O solution and purified using HPLC. The solution phase modification was performed after cleavage and purification of the peptide using HPLC.

Acylated peptides were purified on the Shimadzu LC System with a Phenomenex KinetexTM 5 μm XB-C18 (250 × 21.1 mm, 100 Å) column using different gradients of 0.1 % trifluoroacetic acid (TFA) in H_2_O (solvent A) and 0.1 % TFA in acetonitrile (solvent B) solutions. UPLC-MS analysis was performed using Waters Acquity UPLC-MS system (Milford, USA) with a Waters Acquity-UPLC-MS-BEH C18; 1.7 μM (2.1 × 50 mm; 30 Å) column. As the mobile phase, 0.1 % formic acid in H_2_O (solvent A) and 0.1 % formic acid in acetonitrile (ACN; solvent B) solutions were used. A typical gradient from 95:5 (v/v)H_2_O:ACN to 5:95 (v/v)H_2_O:ACNin 6 mins was used for most of the runs. Data analysis was performed using Waters MassLynx software.

### HPLC-based deacylation assays

Individual Abz-labeled fluorescent peptides (10 µM) were incubated with 100 nM HDAC11 at 37°C for 30 mins. in an assay buffer comprising 50 mM HEPES, 140 mM NaCl, 10 mM KCl, 2 mg/ml BSA, 1 mM TCEP, pH 7.4. The reaction was terminated by the addition of acetic acid to a final concentration of 0.5 % and the reaction products quantified by means of RP-HPLC (Shimadzu, HPLC Prominence system) on a column Kinetex^®^ 2.6 µm XB-C18 100 Å, 100 × 3 mm (Phenomenex, Torrance, CA, USA). The fluorescence detection with excitation/emission wavelengths set at 320/420 nm, respectively, was used together with a calibration curve of known concentrations of the fluorescent reaction product.

### Steady-state kinetics

Individual acylated peptides in the concentration range of 0.01 – 200 µM were incubated with the optimized concentration (1.5 – 100 nM) of HDAC11 at 37°C for 30 mins in an assay buffer. The reaction was quenched by the addition of acetic acid to a final concentration of 0.5 % and the reaction product quantified as above. The data were fitted using the GraphPad Prism software (GraphPad Software, San Diego, CA, USA) and kinetic values calculated through non-linear regression analysis.

### Determination of inhibition constant

The inhibition constants for free fatty acids were determined using the modified HPLC-based assay with trifluoroacetyl peptide as the substrate. Briefly, the tested compounds were preincubated with hsHDAC11 for 10 mins at 37°C and the deacylation reaction started by adding the 50 µM substrate. Following a 30-min incubation at 37°C, the reaction was stopped by the addition of 0.5% acetic acid, and reaction products were quantified using HPLC. The data were fitted using the GraphPad Prism software, and IC_50_ values were calculated by non-linear regression analysis. The inhibitor-free and enzyme-free controls defined 100% and 0% HDAC11 activity, respectively.

### Homology modeling

The amino acid sequence of hsHDAC11 was retrieved from the protein database of NCBI (NP_079103.2) and BLASTed to find a suitable template structure. Among other HDAC structures, the crystal structure of HDAC8 (3MZ4) was selected because: (i) HDAC8 is thus far the only Zn-dependent HDAC for which the fatty-deacylase activity has been described; (ii) the crystal structure was determined to high (1.85 Å) resolution limits. Following the selection of the template, the software Modeller 9.14 was used to construct the target-template sequence alignment and to calculate the 3D homology model of HDAC11. To predict the possible site occupied by FA-acyl substrates, we have performed the docking experiment with decanoylated Gly-[Lys-decanoyl]-Phe tripeptide. The software AutoDockTool 1.5.6 was used for the preparation of ligand and receptor molecules. The docking experiment was completed using AutoDock Vina 1.1.2. The docking area was defined by a cube centered at the catalytic H142 with dimensions 24 Å x 24 Å x 24 Å. The protein was handled as rigid and ligand as flexible during the docking. Binding site visualization and interaction pattern analysis were executed with PyMOL, 0.99rc6.

### SDS-PAGE

Denatured and reduced samples were loaded into 10% polyacrylamide gel and run in Tris-glycine-SDS running buffer at 150 V for 90 mins. The gel was stained with Coomassie Blue G-250 to allow for the visualization of individual proteins.

## ACKNOWLEDGMENTS

We thank to Barbora Havlinova and Petra Baranova for the excellent technical assistance, Dr. Glenda Alquicer for the cloning of hsHDAC11, prof. E. Seto (The George Washington University, D.C., USA) for the original human HDAC11 clone, and K. Harant and P. Talacko (Laboratory of Mass spectrometry, BIOCEV, Charles University) for MS analysis. This work was supported by the Czech Science Foundation (grant No 15-19640S to C.B.), the CAS (RVO: 86652036) and the project „BIOCEV” (CZ.1.05/1.1.00/02.0109) from the ERDF.

## ADDITIONAL INFORMATION

Supplementary Material and Methods, Supplementary figures 1 – 19, Supplementary table 1 (PDF)

### Competing financial interests

The authors declare to have no competing financial interests in relation to the work described within the manuscript file.

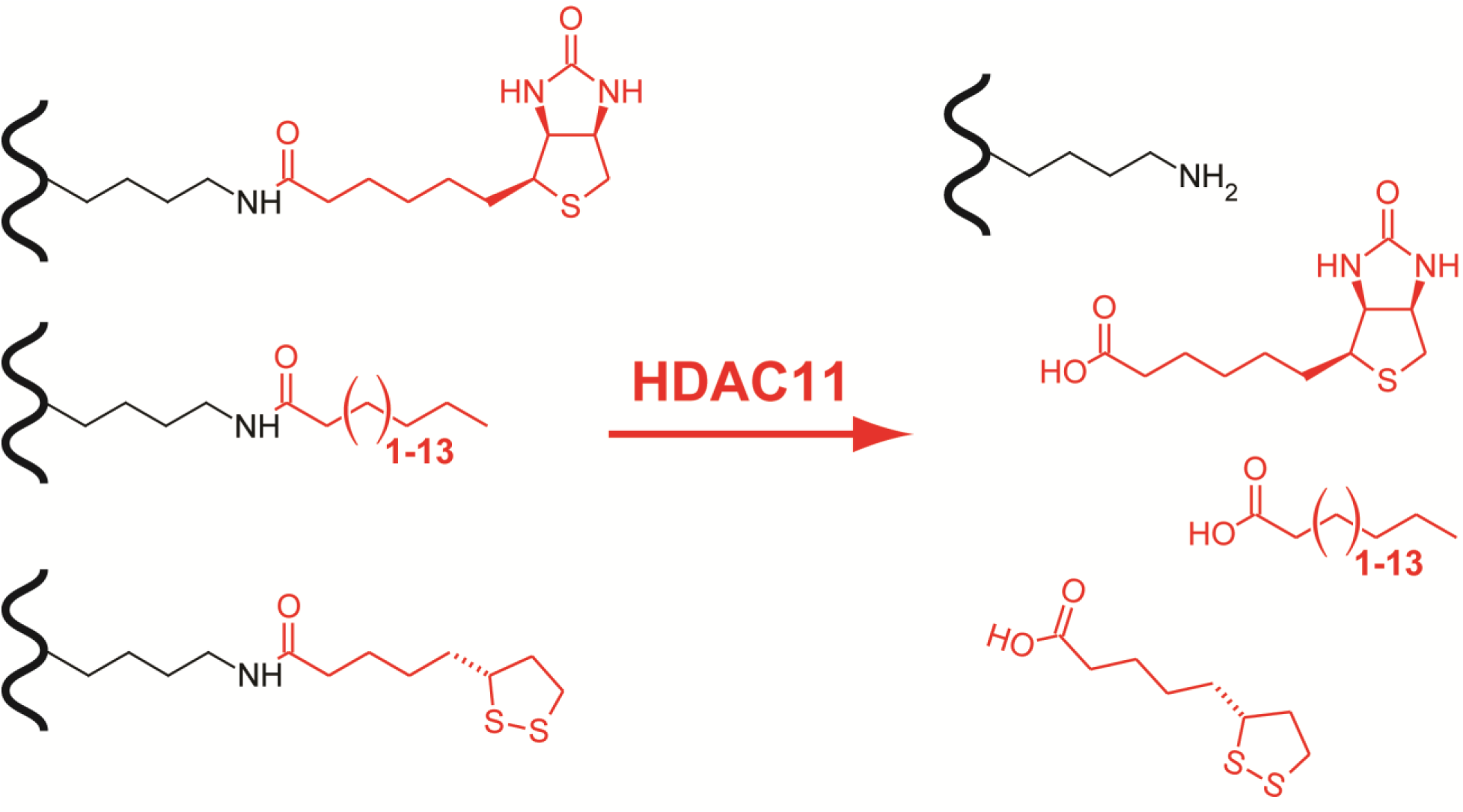

